# Differential physiological, behavioral, and medial prefrontal cortex transcriptomic responses to chronic restraint stress between BALB/c and C57BL/6J mice

**DOI:** 10.64898/2026.08.17.745147

**Authors:** Taiga Kurihara, Aoi William Omi, Yuka Nakasone, Asuka Inami, Teruki Shirayama, Akiko Matsumoto, Ikumi Endo, Goki Yamada, Shiori Kawase, Eiko Kato, Misato Yasumura, Hiroki Yasuda, Takeshi Uemura

## Abstract

Chronic stress is a major risk factor for psychiatric disorders such as depression and anxiety, yet the biological basis of individual differences in stress susceptibility and resilience remains poorly understood. Here, we examined physiological, behavioral, and medial prefrontal cortex (mPFC) transcriptomic responses to chronic restraint stress (CRS) in male BALB/c and C57BL/6J mice. After 21 days of CRS, BALB/c mice exhibited greater stress-related changes than C57BL/6J mice, including greater body weight loss, elevated serum corticosterone, reduced serum antioxidant capacity, and more pronounced depression-like behaviors. RNA sequencing showed largely strain-specific transcriptional changes in the mPFC. Strain × stress interaction analysis, followed by canonical pathway analysis using Ingenuity Pathway Analysis (IPA), identified strain-dependent molecular signatures. The most prominent differences involved extracellular matrix (ECM) organization and remodeling and neuroinflammatory signaling pathways, with greater predicted activation in BALB/c mice. IPA upstream regulator analysis further predicted multiple candidate regulators associated with these pathways, including TGF-β/SMAD, C4a/C4b, and MAPK14. Among genes associated with these pathways, several ECM-related genes were preferentially upregulated in BALB/c mice, whereas activity-dependent immediate early genes were preferentially downregulated in C57BL/6J mice. These findings suggest that the strain-dependent mPFC transcriptional programs identified here may contribute to differential stress susceptibility and resilience.

## Introduction

Chronic stress is a major environmental risk factor for psychiatric disorders, such as major depressive disorder (MDD), anxiety disorders, and post-traumatic stress disorder (PTSD)^1–3^. Prolonged stress exposure induces structural and functional alterations in several brain regions, including the hippocampus, amygdala, and medial prefrontal cortex (mPFC) ^4–7^. These stress-induced neurobiological changes are thought to contribute to the development of psychiatric disorders.

However, exposure to chronic stress does not inevitably result in psychopathology. Behavioral and physiological responses to stress vary considerably among individuals, forming the basis for the concepts of stress susceptibility and resilience^8,9^. Although stress-related psychiatric disorders such as MDD, PTSD, and anxiety disorders are moderately heritable and highly polygenic, genetic factors alone do not determine disease risk^10,11^. Numerous studies suggests that individual differences in stress susceptibility and resilience are shaped by interactions between genetic background and environmental stressors, commonly referred to as gene–environment (G × E) interactions^10,12^. At the molecular level, stress-induced transcriptional changes in stress-responsive brain regions have been implicated in individual differences in stress susceptibility and resilience^13–16^.

Because human studies are often complicated by genetic and environmental variability, animal models are indispensable for elucidating the biological basis of G × E interactions. Various rodent stress paradigms, including chronic restraint stress (CRS), chronic social defeat stress, and chronic unpredictable stress, have been widely used to examine stress-induced behavioral changes and molecular alterations in the brain^17^. Recent transcriptomic studies have shown that stress-induced gene expression changes in the brain differ depending on the stress paradigm and exposure duration^18^.

Among mouse strains, BALB/c and C57BL/6J mice show well-characterized differences in stress-induced behavioral changes and physiological responses, with BALB/c mice generally show more pronounced behavioral and physiological responses to stress than C57BL/6J mice. ^15,19–21^. Consistent with these phenotypic differences, gene expression studies have demonstrated that chronic stress induces distinct, and in some cases opposing, transcriptional responses between these strains^22,23^, indicating that genetic background profoundly influences molecular responses to stress.

Among stress-responsive brain regions, the mPFC is of particular interest because it plays a central role in emotional regulation, cognitive control, and top-down modulation of the hypothalamic–pituitary–adrenal axis^24,25^. Chronic stress induces structural remodeling, dendritic retraction, synaptic alterations, and changes in neuronal activity within the mPFC^3,6,7,26^. Recent studies have further shown that chronic stress induces alterations in medial prefrontal synaptic and circuit mechanisms, as well as transcriptional responses, that are associated with stress-related behavioral phenotypes^15,16,27^. Nevertheless, comprehensive studies integrating physiological, behavioral, and transcriptomic responses to CRS in BALB/c and C57BL/6J mice remain limited. In the present study, we examined physiological, behavioral, and transcriptomic responses to CRS in male BALB/c and C57BL/6J mice. BALB/c mice showed greater physiological responses and more pronounced depression-like behaviors than C57BL/6J mice. Using RNA sequencing of the mPFC, we identified genes showing strain-dependent responses to CRS. Ingenuity Pathway Analysis (IPA) canonical pathway and upstream regulator analyses further predicted strain-dependent changes in extracellular matrix (ECM) organization and remodeling, neuroinflammatory signaling, and activity-dependent transcription, together with candidate upstream regulators.

## Results

### CRS differentially affects body weight changes in BALB/c and C57BL/6J mice

To investigate strain differences in responses to chronic stress, male BALB/c and C57BL/6J mice were subjected to CRS for 6 h per day over 21 consecutive days (Fig. 1A). Body weight was measured daily throughout the experimental period (Fig. 1B, C). Control (CTL) mice in both strains showed a progressive increase in body weight over 21 days, whereas CRS altered the trajectory of body weight change in a strain-dependent manner. In BALB/c mice, CRS suppressed the normal body weight gain observed in CTL mice, with body weight remaining below baseline throughout the experiment (Fig. 1B). A similar suppression of body weight gain was observed in C57BL/6J mice. Unlike BALB/c mice, however, body weight remained close to baseline during the early phase of CRS and gradually increased after day 14, although the difference between day 1 and day 21 did not reach statistical significance (*p* = 0.059) (Fig. 1C). These results suggested that CRS has a greater impact on body weight in BALB/c mice than in C57BL/6J mice.

**Figure 1.**
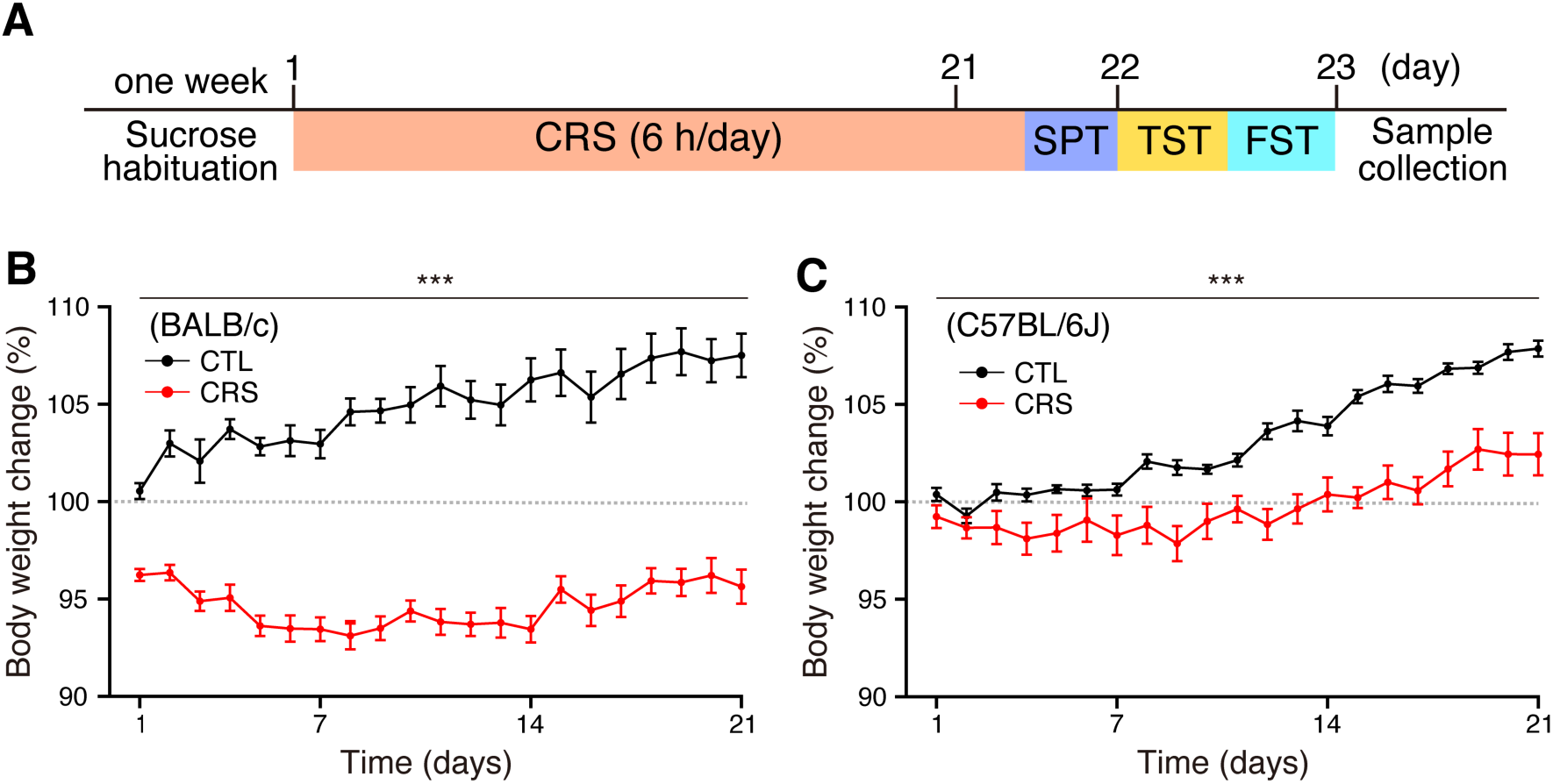
Experimental timeline and strain-dependent effects of CRS on body weight. (A) Schematic overview of the experimental design. Male BALB/c and C57BL/6J mice (9–10 weeks old) were subjected to CRS for 6 h/day over 21 consecutive days. Mice were habituated to sucrose solution for 1 week before CRS. The sucrose preference test (SPT) was performed on day 21, followed by the tail suspension test (TST) and forced swim test (FST) on day 22. On day 23, brain tissue, adrenal glands, and blood samples were collected for subsequent analyses. (B, C) Changes in body weight during CRS. Body weight changes in (B) BALB/c mice (CTL, n = 10; CRS, n = 11) and (C) C57BL/6J mice (CTL, n = 10; CRS, n = 8) are expressed as a percentage relative to the initial body weight measured before CRS. Body weight changes from 1 day after the initiation of CRS (day 1) through day 21 are shown. Data are presented as mean ± SEM. Statistical analyses for time-course changes were performed using a two-way repeated measures ANOVA [BALB/c: group × time interaction, *F*(20, 380) = 8.92, *p* < 0.0001; C57BL/6J: group × time interaction, *F*(20, 320) = 10.4, *p* < 0.0001]. *** *p* < 0.001 for the group × time interaction.

### CRS induces physiological stress responses in both strains

To assess physiological stress responses, adrenal gland weight, serum corticosterone levels, and serum oxidative stress markers were measured after the completion of CRS and the subsequent behavioral tests (Fig. 1A). CRS significantly increased adrenal gland weight in both BALB/c and C57BL/6J mice compared with their respective controls (Fig. 2A). Serum corticosterone levels were significantly elevated in BALB/c mice following CRS, whereas no significant change was detected in C57BL/6J mice (Fig. 2B). To evaluate oxidative stress status, serum biological antioxidant potential (BAP) and diacron-reactive oxygen metabolite (d-ROM) levels were measured. BAP levels were significantly reduced in BALB/c mice following CRS but remained unchanged in C57BL/6J mice (Fig. 2C). In contrast, d-ROM levels were significantly increased in both strains following CRS (Fig. 2D). These results suggested that BALB/c mice exhibit stronger endocrine response and more pronounced disruption of oxidative status following CRS than C57BL/6J mice.

**Figure 2.**
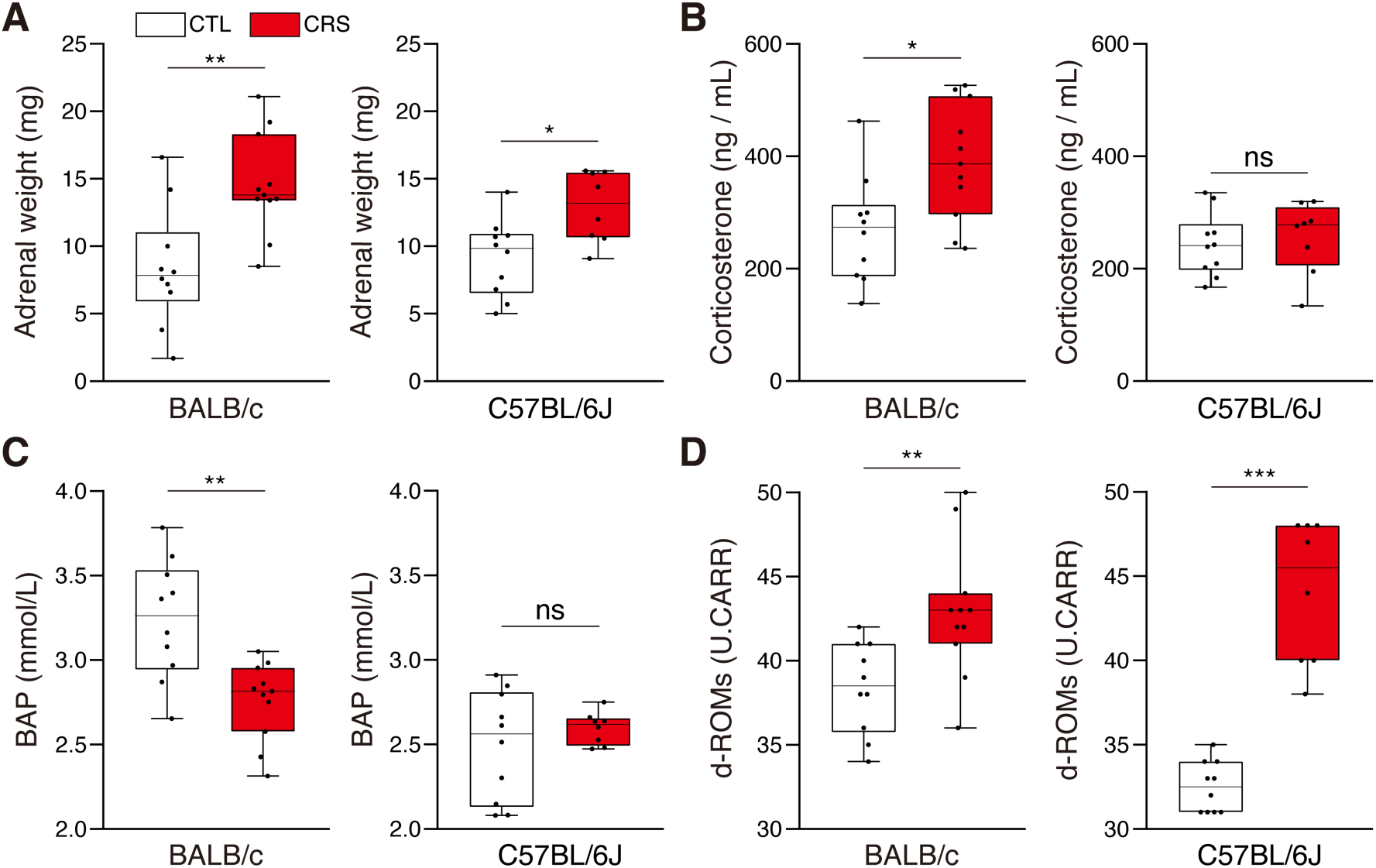
Effects of CRS on adrenal gland weight, serum corticosterone levels, and serum oxidative stress markers. (A) Bilateral adrenal gland weights in BALB/c and C57BL/6J mice measured 2 days after the final CRS session (6 h/day for 21 consecutive days). (B–D) Blood samples were collected from BALB/c and C57BL/6J mice 2 days after the final CRS session, and serum levels of corticosterone (B), BAP (C), and d-ROMs (D) were measured. The data are presented as boxplots (BALB/c mice: CTL, n = 10; CRS, n = 11; C57BL/6J mice: CTL, n = 10; CRS, n = 8). The horizontal line in each box shows the median; the box shows the interquartile range (IQR); and the whiskers represent 1.5 × IQR. BAP, biological antioxidant potential; d-ROMs, reactive oxygen metabolites; ns, not significant; U.CARR, Carratelli units. \*\*\**p* < 0.001, \*\**p* < 0.01, \**p* < 0.05; Mann–Whitney U test.

### BALB/c mice show greater susceptibility to CRS-induced depression-like behaviors

Next, we examined the effects of 21-day CRS on depression-like behaviors using the sucrose preference test (SPT), tail suspension test (TST), and forced swim test (FST) (Fig. 3). In the SPT, CRS significantly reduced sucrose preference in BALB/c mice, whereas no significant change was observed in C57BL/6J mice (Fig. 3A), indicating anhedonia-like behavior in BALB/c mice. In the TST and FST, which assess despair-like behavior, immobility time was significantly increased in BALB/c mice following CRS (Fig. 3B, C). In contrast, C57BL/6J mice showed a significant increase in immobility time only in the FST, with no significant change in the TST (Fig. 3B, C). Together, these results suggested that BALB/c mice are more susceptible to CRS-induced depression-like behaviors than C57BL/6J mice.

**Figure 3.**
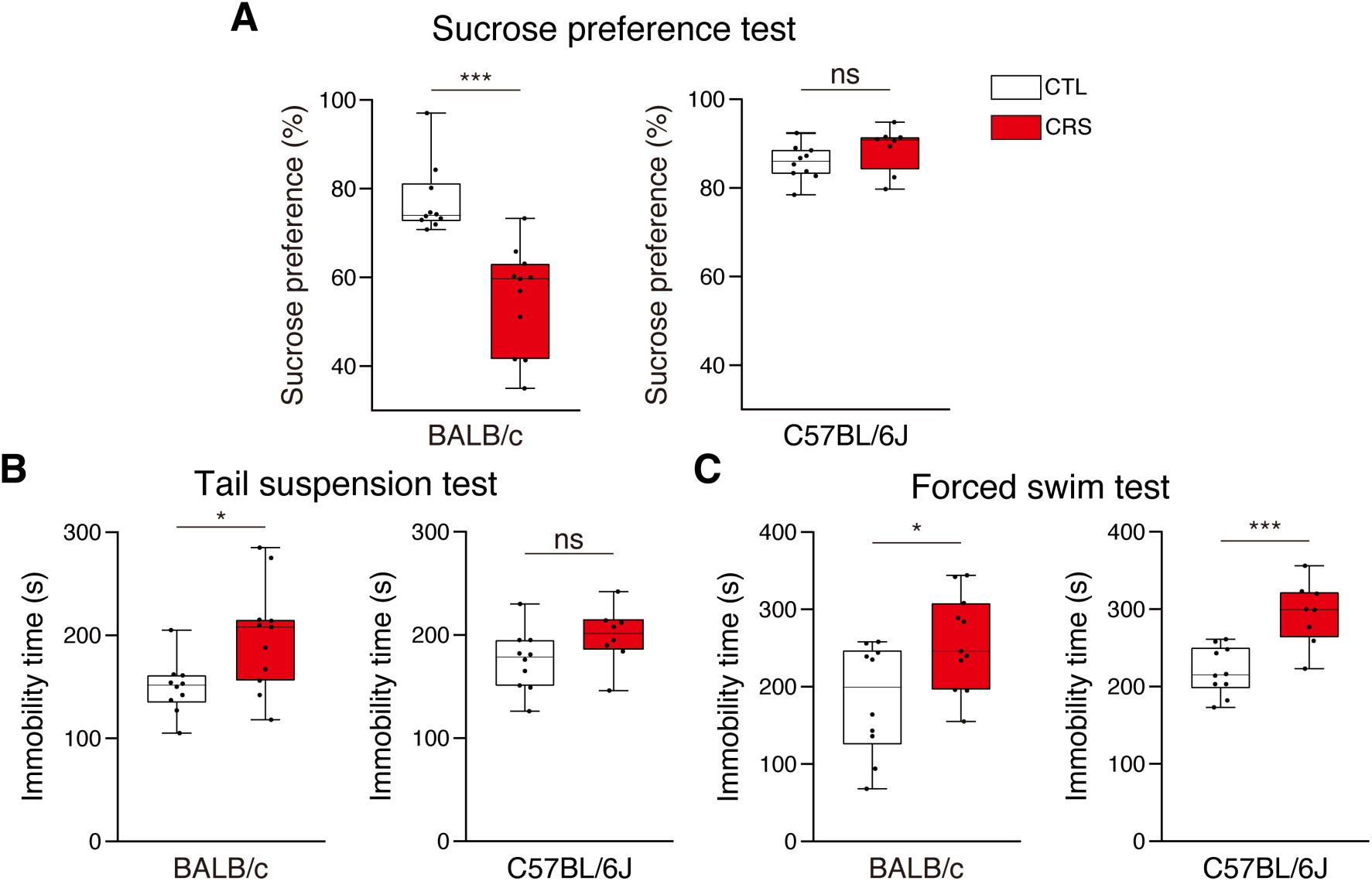
Effects of CRS on depression-like behaviors in BALB/c and C57BL/6J mice. (A–C) The effects of CRS (CRS; 6 h/day for 21 consecutive days) were assessed using the sucrose preference test (A), tail suspension test (B), and forced swim test (C) in BALB/c and C57BL/6J mice. The data are presented as boxplots (BALB/c mice: CTL, n = 10; CRS, n = 11; C57BL/6J mice: CTL, n = 10; CRS, n = 8). The horizontal line in each box shows the median; the box shows the IQR; and the whiskers represent 1.5 × IQR. \*\*\**p* < 0.001, \**p* < 0.05; Mann–Whitney U-test. ns, not significant.

### CRS induces distinct transcriptomic responses in the mPFC of BALB/c and C57BL/6J mice

To investigate molecular responses to CRS, RNA sequencing was performed using mPFC samples collected after behavioral testing (Fig. 1A). Differential expression analysis identified 591 differentially expressed genes (DEGs) in BALB/c mice (239 downregulated and 352 upregulated) and 568 DEGs in C57BL/6J mice (274 downregulated and 294 upregulated) following CRS (Fig. 4A, B). Of these, 477 DEGs were unique to BALB/c mice, 454 were unique to C57BL/6J mice, and only 114 were shared between the two strains (Fig. 4C), indicating that the majority of stress-responsive transcriptional changes were strain-specific.

**Figure 4.**
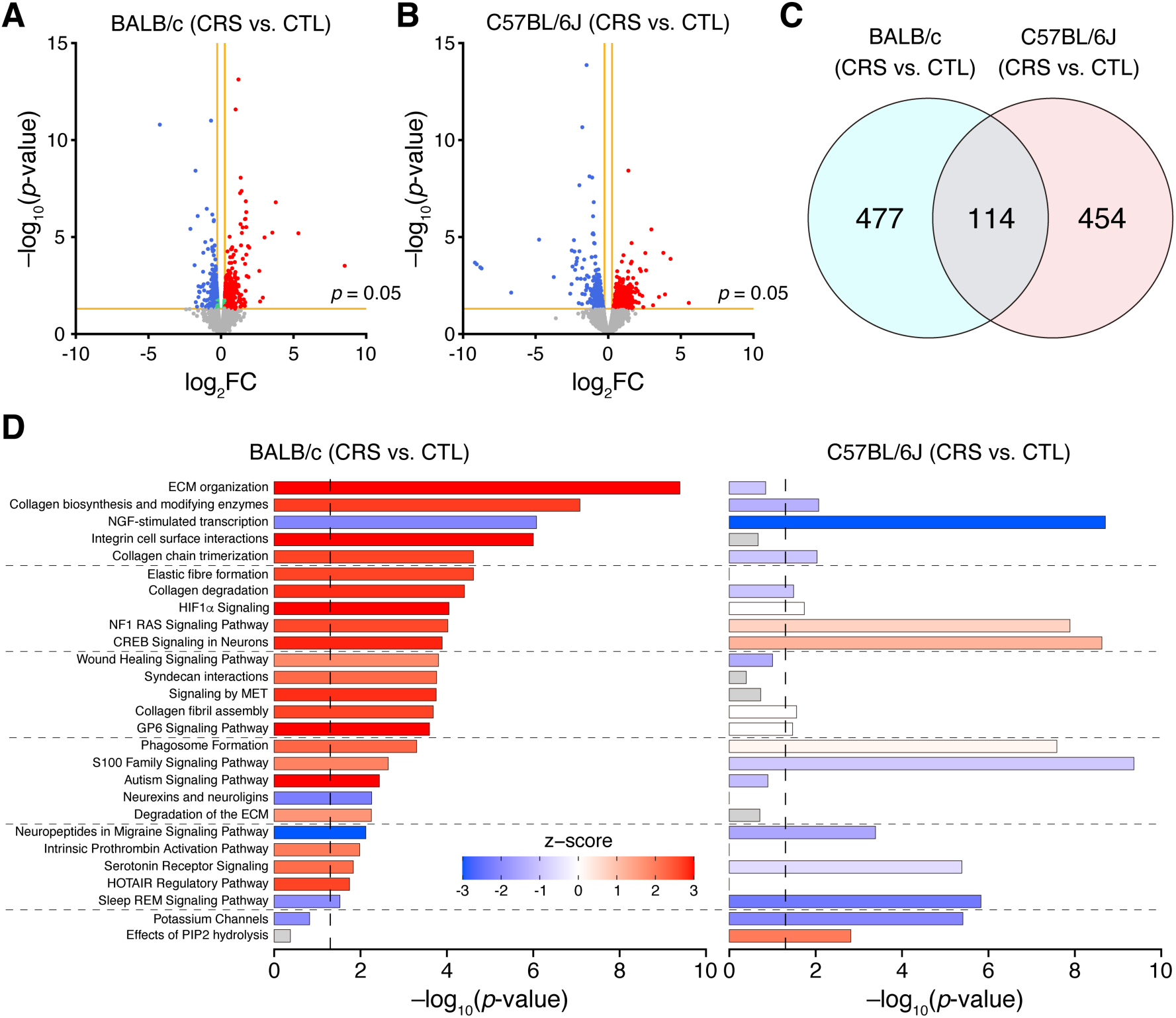
Strain-specific transcriptomic responses and canonical pathway alterations induced by CRS. (A, B) Volcano plots of differentially expressed genes (DEGs) in the mPFC of BALB/c (A) and C57BL/6J (B) mice following CRS. The vertical orange lines indicate the threshold for |log_2_FC| > 0.263, and the horizontal orange line indicates the threshold for *p* < 0.05. (C) Venn diagram showing the numbers and overlap of DEGs identified in BALB/c and C57BL/6J mice. (D) IPA canonical pathway analysis of DEGs identified in BALB/c and C57BL/6J mice following CRS. Representative IPA canonical pathways associated with neuronal signaling, extracellular matrix (ECM) organization, and immune/inflammatory responses are shown. All pathway names shown correspond to canonical pathways as defined in IPA. For clarity, “Collagen fibril assembly” is a shortened form of the full IPA pathway name “Assembly of collagen fibrils and other multimeric structures”. Bar length represents the −log_10_(*p*-value), and the vertical dashed line indicates the significance threshold (*p* = 0.05). Bar colors represent the IPA activation z-score (red, predicted activation; blue, predicted inhibition; gray, no prediction).

To explore the biological significance of these DEGs, Gene Ontology (GO) and pathway enrichment analyses were performed using Metascape (Supplementary Fig. 1A, B). QIAGEN IPA canonical pathway analysis further predicted distinct patterns of canonical pathway activation and inhibition between the two strains (Fig. 4D, Supplementary Fig. 2A, B). In BALB/c mice, several IPA-defined ECM-related pathways consistently showed positive IPA activation z-scores following CRS, indicating predicted pathway activation. These included ECM organization, collagen biosynthesis and modifying enzymes, and integrin cell surface interactions. Similarly, IPA-defined pathways in the cellular stress and injury category, including hypoxia-inducible factor 1α (HIF1α) signaling, wound healing signaling, and S100 family signaling, also showed positive IPA activation z-scores in BALB/c mice. Because IPA canonical pathways are defined across diverse tissue types, those with names related to fibrosis or wound healing should be interpreted in the brain as reflecting shared molecular programs of ECM remodeling and stress response rather than fibrosis itself. In contrast, in C57BL/6J mice, pathways in both the ECM organization and cellular stress and injury categories showed negative or near-zero IPA activation z-scores, indicating predicted inhibition or no clear prediction of activation of these pathways following CRS. These opposing patterns of predicted canonical pathway activity suggested distinct molecular responses to CRS between the two strains.

### Interaction analysis reveals strain-dependent transcriptional changes and IPA-predicted pathway activity

These opposing patterns of predicted pathway activity prompted a direct comparison of strain-dependent transcriptional responses using a strain × stress interaction analysis. The interaction log₂ fold change (log₂FC) was defined as the difference in CRS-induced transcriptional changes between BALB/c and C57BL/6J mice (Fig. 5A). Interaction analysis identified 302 genes with significant interaction effects, of which 174 showed negative interaction effects and 128 showed positive interaction effects (Fig. 5B). Negative log₂FC values indicated relatively greater transcriptional responses in BALB/c mice, whereas positive values indicated relatively greater responses in C57BL/6J mice.

**Figure 5.**
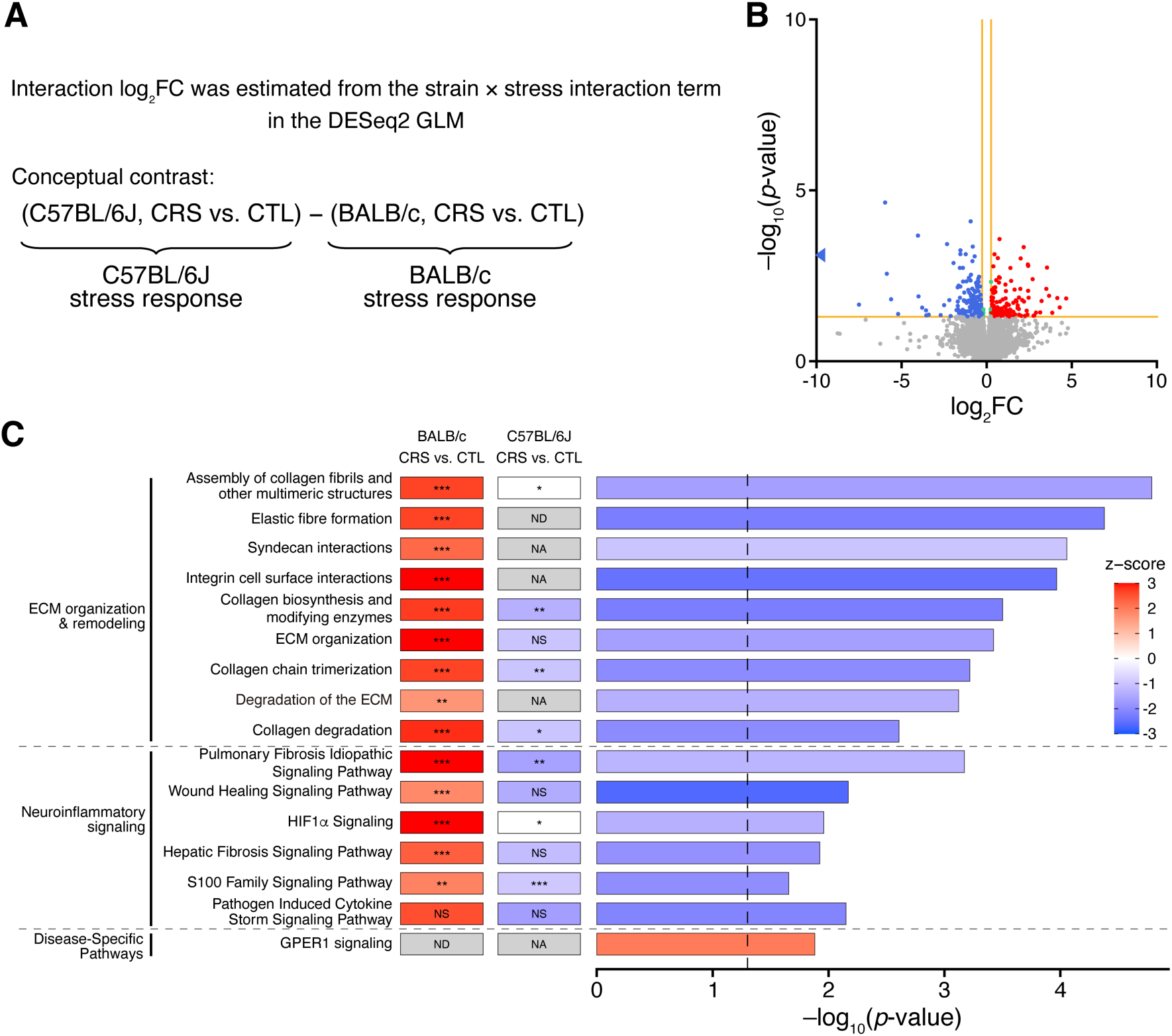
Differentially expressed genes and predicted pathway activity based on strain × stress interaction analysis. (A) Definition of interaction log₂FC values. Interaction log₂FC values were estimated from the strain × stress interaction term in DESeq2. Negative values indicate a smaller CRS-induced log₂FC in C57BL/6J mice than in BALB/c mice, whereas positive values indicate a larger CRS-induced log₂FC in C57BL/6J mice. (B) Volcano plot showing differentially expressed genes (DEGs) identified by the strain × stress interaction analysis. The vertical and horizontal orange lines indicate the thresholds of |log_2_FC| > 0.263, and *p* < 0.05, respectively. Red and blue dots represent genes with significant positive and negative interaction effects, respectively. The blue triangle indicates a transcript with an interaction log₂FC less than −10, which lies outside the plotted x-axis range. (C) IPA canonical pathway analysis based on interaction DEGs. Representative IPA canonical pathways associated with stress responses are shown. Pathways were selected based on their statistical significance and/or activation z-scores. The two columns on the left show the significance of the corresponding pathways in the within-strain comparisons (CRS vs. CTL) for BALB/c and C57BL/6J mice (\**p* < 0.05, \*\**p* < 0.01, \*\*\**p* < 0.001; ND, no data; NA, not available; NS, not significant). The labels on the left indicate functional categories. Bar length represents the −log_10_(*p*-value), and the vertical dashed line indicates the significance threshold (*p* = 0.05). Bar colors represent the IPA activation z-score (red, predicted activation; blue, predicted inhibition). For the interaction analysis, negative IPA z-scores indicate relatively lower predicted activation in C57BL/6J mice, or conversely, greater predicted activation in BALB/c mice. Positive IPA z-scores indicate the opposite pattern. GLM, generalized linear model.

To characterize the biological functions represented by the interaction DEGs, GO and pathway enrichment analyses were performed using Metascape (Supplementary Fig. 1C). IPA canonical pathway analysis further predicted distinct pathway activity profiles between the two strains (Fig. 5C and Supplementary Fig. 2C). Canonical pathways were grouped according to IPA classifications and further summarized into broader functional categories for interpretation. Pathways classified by IPA under ECM organization, including assembly of collagen fibrils, elastic fibre formation, integrin cell surface interactions, and collagen biosynthesis and modifying enzymes, were among the most statistically significant pathways and consistently showed negative IPA activation z-scores in the interaction analysis; hereafter, we refer to these pathways collectively as ECM organization and remodeling. Given that the interaction contrast was defined as the C57BL/6J stress response minus the BALB/c stress response, these negative z-scores indicate that IPA predicted relatively greater activation of these pathways in BALB/c mice than in C57BL/6J mice. Pathways classified by IPA under cellular stress and injury, including wound healing signaling and HIF1α signaling, showed similar negative z-scores. The pathogen-induced cytokine storm signaling pathway, classified by IPA under signal transduction and not significantly enriched in either strain, also showed a negative interaction z-score. Because these injury- and immune-responsive pathways are, in the brain, closely associated with neuroinflammatory processes, we hereafter refer to them collectively as neuroinflammatory signaling. Unlike these pathways, GPER (G protein-coupled estrogen receptor) 1 signaling, classified by IPA under disease-specific pathways, showed a positive IPA activation z-score, suggesting relatively greater predicted activation in C57BL/6J mice. Together, these results suggested distinct pathway alterations associated with strain-dependent transcriptional responses to CRS.

### Upstream regulator analysis predicts candidate regulators associated with strain-dependent stress responses

To infer candidate upstream regulators associated with the strain-dependent transcriptional differences, interaction DEGs were analyzed using the IPA upstream regulator analysis tool (Fig. 6A). For visualization, representative predicted upstream regulators related to the major canonical pathway categories were selected and grouped broadly into three functional categories based on their biological roles: ECM organization and remodeling, neuroinflammatory signaling, and neural activity-related signaling. Predicted regulators associated with ECM organization and remodeling included members of the TGF-β family (TGFB1, TGFB2, and TGFB3), TGFBR2, SMAD3, FKBP10, PEAR1, SRF, ALPHA CATENIN family, miR-29b-3p, and miR-21-5p. The remaining regulators were grouped as neuroinflammatory signaling (C4a/C4b and MAPK14) or neural activity-related signaling (dopamine, APLNR, and NGEF).

**Figure 6.**
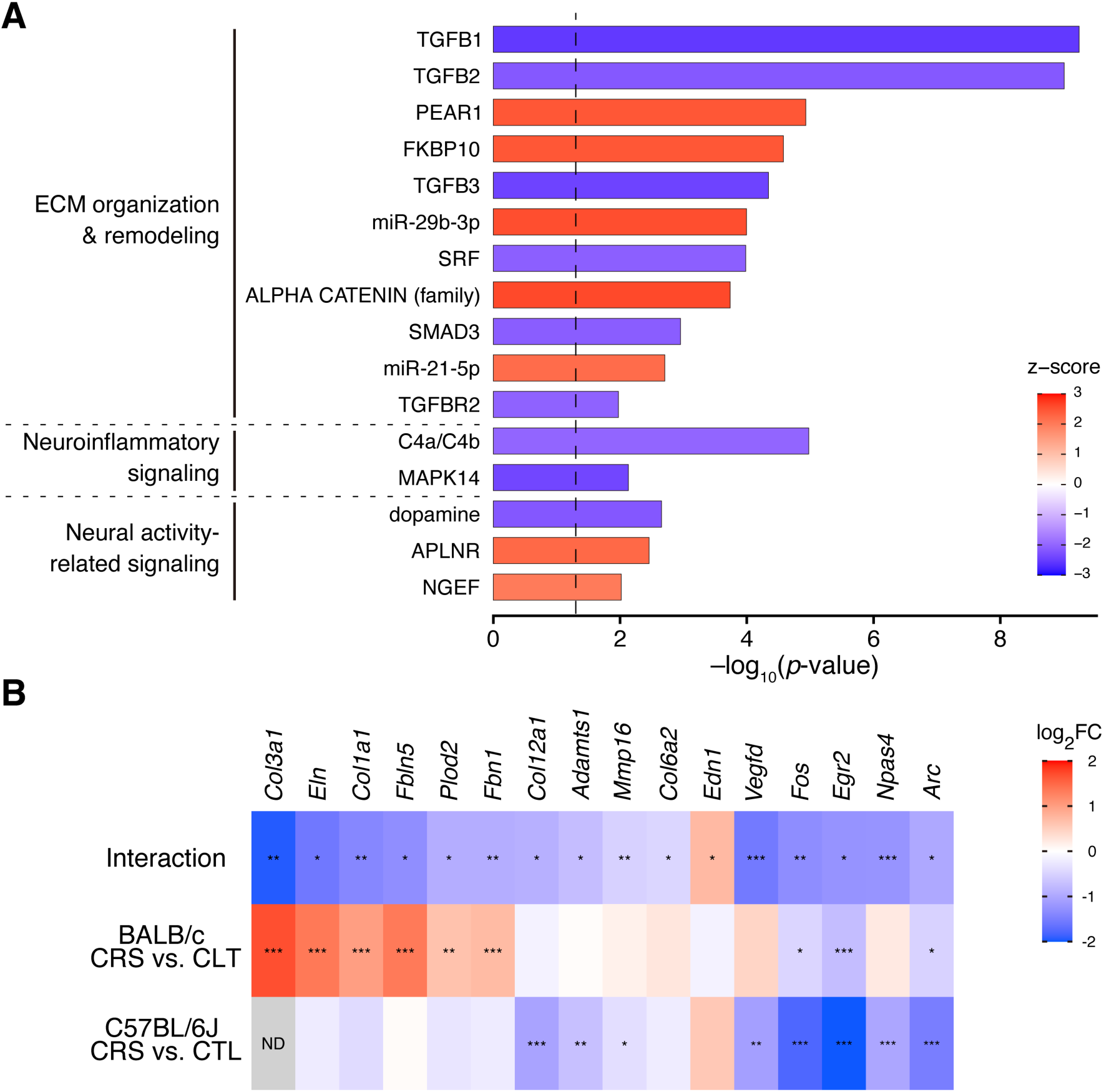
Predicted upstream regulators and downstream target genes identified by interaction analysis. (A) Upstream regulators predicted by IPA based on the interaction DEGs defined by (C57BL/6J CRS vs. CTL) − (BALB/c CRS vs. CTL). Representative predicted upstream regulators associated with the IPA canonical pathways identified from the interaction DEGs are shown. Representative upstream regulators were selected for visualization based on statistical significance, biological interpretability, relevance to the major canonical pathway categories, and an absolute IPA activation z-score ≥2. Exogenous chemicals, drugs, toxicants, and reagent compounds were excluded unless directly relevant. Regulators are grouped into three functional categories: ECM organization and remodeling, neuroinflammatory signaling, and neural activity-related signaling. Bar length represents the −log_10_(*p*-value), and the vertical dashed line indicates the significance threshold (*p* = 0.05). Bar colors represent the IPA activation z-score (red, predicted activation; blue, predicted inhibition). For the interaction analysis, negative z-scores indicate relatively lower predicted activation in C57BL/6J mice, or conversely relatively greater predicted activation in BALB/c mice. Positive z-scores indicate the opposite pattern. (B) Heatmap showing representative IPA-defined downstream target genes associated with the predicted upstream regulators. Columns represent log_2_FC values derived from the interaction analysis (top), the BALB/c CRS vs. CTL comparison (middle), and the C57BL/6J CRS vs. CTL comparison (bottom). Colors represent the direction and magnitude of log₂FC values (red, positive; blue, negative). \*\*\**p* < 0.001, \*\**p* < 0.01, and \**p* < 0.05; ND, no data.

To examine transcriptional changes in downstream target genes associated with these predicted regulators, we focused on representative genes defined by IPA annotations (Fig. 6B and Supplementary Table 1). The interaction analysis revealed that many ECM-related genes, including *Col3a1*, *Eln*, *Col1a1*, *Fbln5*, *Plod2*, *Fbn1*, *Col12a1*, *Adamts1*, *Mmp16*, and *Col6a2*, showed strain-dependent CRS responses. Among these genes, some interactions reflected stronger upregulation in BALB/c mice, whereas others reflected downregulation in C57BL/6J mice. These genes were linked in IPA to predicted regulators such as the TGF-β family, SMAD3, PEAR1, FKBP10, and miR-29b-3p. In contrast, activity-dependent immediate early genes (IEGs), including *Fos*, *Egr2*, *Npas4*, and *Arc*, were more strongly downregulated by CRS in C57BL/6J mice. In the IPA analysis, these genes were listed as target genes of multiple predicted regulators spanning two categories, including C4a/C4b and dopamine, as well as TGF-β family members (TGFB1, TGFB2, and TGFB3) and their downstream effector SMAD3 (Supplementary Table 1). Vascular-regulatory genes, including *Edn1* and *Vegfd*, were also linked to predicted upstream regulators such as the TGF-β family members and SRF.

## Discussion

The present study demonstrates that male BALB/c and C57BL/6J mice exhibit markedly different responses to CRS at physiological, behavioral, and transcriptomic levels. Relative to C57BL/6J mice, BALB/c mice showed greater body weight loss, elevated corticosterone levels, reduced antioxidant capacity, and more pronounced depression-like behavioral abnormalities (Figs. 1–3). RNA sequencing of the mPFC further revealed that these divergent phenotypic outcomes were accompanied by largely strain-specific transcriptional responses to CRS (Fig. 4). Strain × stress interaction analysis identified genes showing strain-dependent stress responses, and IPA analyses predicted strain-dependent changes in biological pathways, including ECM organization and remodeling, neuroinflammatory signaling, and neural activity-dependent transcription (Figs. 5 and 6). Together, these findings provide a comprehensive view of physiological, behavioral, and molecular features associated with differential susceptibility and resilience to CRS.

The pronounced physiological and behavioral sensitivity of BALB/c mice to CRS observed here (Figs. 1–3) is consistent with previous studies showing that BALB/c mice are more susceptible to stress than C57BL/6J mice, with greater corticosterone responses and more severe depression-like behaviors^15,19,20^. Both strains showed increased d-ROM levels following CRS, indicating elevated oxidative stress. However, only BALB/c mice showed reduced BAP levels, suggesting that their antioxidant capacity was more easily impaired by chronic stress. A previous study reported that chronic stress increased reactive oxygen species levels in the hippocampus of BALB/c mice but not C57BL/6J mice^28^. Although the present study differs from that report in that d-ROM levels increased in both strains, the overall pattern is consistent, because the strain-specific reduction in BAP indicates that BALB/c mice are more vulnerable to oxidative stress. This difference may partly reflect the biological samples analyzed, as we measured serum oxidative markers (Fig. 2), whereas reactive oxygen species were measured in the hippocampus in the previous study^28^. Taken together, these findings suggest that altered oxidative status may contribute to the higher stress susceptibility of BALB/c mice.

BALB/c mice showed reduced sucrose preference and increased tail suspension immobility following CRS, whereas forced swim immobility was elevated in both strains (Fig. 3). The selective reduction in sucrose preference in BALB/c mice suggests that anhedonia-like behavior was more closely associated with the stress-susceptible BALB/c phenotype. To identify molecular mechanisms that may underlie these strain-dependent behavioral differences, we focused on genes exhibiting significant strain × stress interactions in the mPFC.

In the interaction canonical pathway analysis, pathways in the ECM organization category, including assembly of collagen fibrils, elastic fibre formation, and integrin cell surface interactions, were among the most statistically significant pathways and were consistently predicted to be more activated in BALB/c mice than in C57BL/6J mice (Fig. 5C). Upstream regulator analysis further predicted members of the TGF-β family (TGFB1–3) and their downstream effector SMAD3 as candidate regulators associated with these ECM-related changes (Fig. 6A). Several ECM-related genes contributing to these IPA-predicted pathway and upstream regulator results showed strain-dependent CRS responses (Fig. 6B). These included genes encoding fibrillar and network-forming collagens (*Col1a1*, *Col12a1*, *Col3a1*, and *Col6a2*), elastic fibre–related genes (*Eln*, *Fbn1*, and *Fbln5*), and ECM-modifying or remodeling enzymes (*Plod2*, *Adamts1*, and *Mmp16*). For approximately half of these genes, the strain-dependent response reflected upregulation in BALB/c mice, whereas for the remainder it reflected downregulation in C57BL/6J mice. Thus, the IPA results suggest that CRS may alter ECM organization and remodeling more strongly in the mPFC of BALB/c mice than in C57BL/6J mice. This interpretation is biologically plausible because TGF-β/SMAD signaling is a well-established regulator of ECM gene expression and tissue remodeling^29^. Moreover, collagen is a major structural component of the brain ECM and is involved in cell-cell interactions among neurons, glia, and vascular cells^30^. Alterations in ECM components and perineuronal nets in the brain are thought to contribute to stress-related neural plasticity and depression-related behavioral and cognitive phenotypes^31–34^. ECM-related transcriptional changes have also been reported in other chronic stress models, such as collagen gene upregulation in the rat hippocampus after chronic immobilization stress and brain region-dependent collagen gene regulation in stress-susceptible mice exposed to chronic agonistic interactions^35,36^. Our findings further show that ECM-related transcriptional responses to CRS differ between BALB/c and C57BL/6J mice in the mPFC.

The ECM has been reported to contribute to vascular and perivascular integrity^30^. Thus, the ECM-related changes observed in the present study may also have implications for BBB function. Interestingly, vascular-regulatory genes such as *Edn1* and *Vegfd*^37,38^ showed strain-dependent CRS responses (Fig. 6B). Previous study has reported that VEGF signaling is involved in stress-induced BBB hyperpermeability and depression-like behaviors in BALB/c mice^9^. However, because BBB integrity was not directly assessed in the present study, this possibility remains to be tested.

In addition to ECM organization and remodeling, pathways related to neuroinflammatory signaling were predicted to be more activated in BALB/c mice than in C57BL/6J mice (Fig. 5C), and upstream regulator analysis further predicted C4a/C4b and MAPK14 as candidate regulators associated with these responses (Fig. 6A). This prediction is consistent with previous report that chronic stress induces neuroinflammatory responses, including microglial activation^40^. In contrast, the weaker predicted ECM- and inflammation-related responses in C57BL/6J mice may be associated with the relatively stress-resilient phenotype of this strain. IPA also showed that several activity-dependent IEGs, such as *Fos*, *Arc*, *Egr2*, and *Npas4*, were shared downstream targets of predicted regulators spanning ECM organization and remodeling, neuroinflammatory signaling, and neural activity-related signaling, including C4a/C4b, dopamine, TGF-β family members, SMAD3, and SRF (Fig. 6B and Supplementary Table 1). These IEGs showed significant strain × stress interactions, with expression reduced more strongly in C57BL/6J mice than in BALB/c mice (Fig. 6B). These findings suggest that altered activity-dependent transcription may be linked to the predicted ECM- and inflammation-associated responses in the mPFC.

These IEGs are rapidly induced by neuronal activity and take part in synaptic plasticity and activity-dependent transcription^41–43^. Among these genes, *Npas4* is notable because mPFC-specific knockdown of *Npas4* has been shown to prevent anhedonia-like behavior in mice exposed to chronic social defeat stress, without affecting social avoidance or anxiety-like behavior^27^. Our data do not allow us to determine whether the coordinated downregulation of these IEGs in C57BL/6J mice reflects lower neuronal activity or an active limit on activity-dependent transcription. The coordinated downregulation of multiple activity-dependent genes raises the possibility that attenuated activity-dependent transcriptional responses contribute to stress resilience in C57BL/6J mice. A previous study showed that chronic chemogenetic activation of somatostatin-expressing (SST) interneurons in the mPFC promoted resilience to chronic variable stress. Interestingly, this behavioral effect of activating mPFC SST neurons was observed only in male mice^44^.

IPA-based analyses also suggested additional candidate pathways and regulators that may be associated with the relatively stress-resilient phenotype of C57BL/6J mice. In particular, GPER1 signaling was predicted to be relatively more activated in C57BL/6J mice (Fig. 5C). Because GPER activation has been reported to exert neuroprotective and anti-inflammatory effects in the brain^45^, this pathway may represent another candidate mechanism associated with the relatively resilient phenotype of C57BL/6J mice. TET1 also showed a near-threshold negative activation z-score in the IPA analysis (Supplementary Table 1). Because Tet1-deficient mice have been reported to show resistance to CRS^14^, TET1-related regulatory mechanisms may warrant further investigation in the context of strain-dependent stress resilience. Although these IPA-based predictions require experimental validation, they provide plausible molecular hypotheses for the strain-dependent responses to chronic stress.

The present study was limited to male mice, and therefore it remains unclear whether the physiological, behavioral, and mPFC transcriptomic differences observed here also occur in females. Given the well-established sex differences in stress responses^46^, future studies including female mice, as well as transcriptomic analyses of other brain regions such as the hippocampus, amygdala, and nucleus accumbens, will be important.

In conclusion, BALB/c and C57BL/6J mice exhibited markedly different physiological, behavioral, and prefrontal transcriptomic responses to CRS. Our findings suggest that stress susceptibility in BALB/c mice is associated with greater IPA-predicted activation of pathways related to ECM organization and remodeling and neuroinflammatory signaling in the mPFC. In contrast, the relatively stress-resilient phenotype of C57BL/6J mice is associated with weaker predicted ECM- and inflammation-related responses and stronger downregulation of activity-dependent IEGs than in BALB/c mice. These findings provide new insight into the molecular basis of strain-dependent responses to chronic stress and highlight potential molecular pathways relevant to stress-related psychiatric disorders.

## Materials and methods

### Approval for animal experiments

The animal care and experimental protocols were reviewed by the Committee for Animal Experiments and approved by the president of Shinshu University (Approval No. 025077) and by the president of Saga University (Approval No. A2022-037-3). All experiments were conducted in accordance with the Guidelines for the Care and Use of Laboratory Animals of Shinshu University and Saga University.

### Animals

Male C57BL/6JmsSlc and BALB/cCrSlc mice were obtained from SLC Japan, Inc. (Tokyo, Japan). Mice were housed in a temperature- and humidity-controlled animal facility (approximately 24°C and 40–60% humidity) under a 12-h light/dark cycle, with 3–5 animals per cage. Food and water were provided ad libitum. After a 1-week acclimation period, male mice aged 9–10 weeks at the start of the experiments were used. After the behavioral tests, all mice were deeply anesthetized by intraperitoneal injection of a mixture of medetomidine hydrochloride, midazolam, and butorphanol tartrate at doses of 0.75, 4.0, and 5.0 mg/kg body weight, respectively. Blood was collected by cardiac puncture, after which the mice were briefly perfused transcardially with physiological saline to remove residual blood. The mice were then euthanized by decapitation, and the brains and adrenal glands were collected.

### Chronic restraint stress model

Mice were restrained in custom-made polyethylene restrainers based on a commercially available disposable restraint cone (Bio Research Center Co., Ltd., Nagoya, Japan). Restraint stress was applied for 6 h per day for 21 consecutive days in the home cage. Control mice were handled only and maintained in their home cages for the same 21-day period. In both the stress and control groups, food and water were withheld during the 6-h period. Body weight was measured daily during CRS.

### Adrenal gland weights and serum biochemical analyses

Two days after the final restraint session (day 23), blood samples were collected from the heart under deep anesthesia, and serum was separated by centrifugation. The serum samples were stored at −80°C until use. After blood collection, both adrenal glands were removed and weighed.

Serum oxidative stress markers (d-ROMs and BAP) were measured using d-ROMs and BAP test reagents (Wismerll Co., Ltd., Tokyo, Japan) with a small-scale, semi-automated 384-well plate method incorporating a pipetting robot, as previously described^47^. The d-ROM values are expressed in Carratelli units (U.CARR).

Serum corticosterone levels were measured using liquid chromatography–tandem mass spectrometry (LC–MS/MS) after solid-phase extraction, as described previously with a slight modification^48^. Briefly, LC-MS grade water was used as eluent A instead of 18 mM acetic acid-ammonium acetate buffer solution at pH 5.3. Serum samples stored at −80 °C were thawed immediately prior to analysis. An internal standard was added to each sample, and then corticosterone was extracted using a C18 solid-phase extraction column (MonoSpin C18; GL Science, Tokyo, Japan). After washing, the compounds were eluted with methanol and mixed with ammonium acetate buffer (buffer:methanol = 6:4, v:v). LC was performed using a C18 analytical column. Corticosterone levels were normalized to the internal standard, corticosterone-d4.

### Sucrose preference test

For sucrose habituation, mice were group-housed (3–5 mice per cage) for 7 days before the CRS. During this period, mice were given free access to two bottles containing 2% sucrose solution (w/v) and tap water, respectively. To avoid side preference in drinking behavior, the positions of the two bottles were switched every day. After CRS (day 21), mice were individually housed and allowed free access to two bottles containing 2% sucrose solution and tap water for 24 h. The intake of sucrose solution and water was measured. Sucrose preference was calculated using the following formula: Sucrose preference (%) = sucrose intake (g) / [sucrose intake (g) + water intake (g)] × 100.

All behavioral tests described below were performed after the mice had been allowed to adapt to the experimental room for at least 1 h before testing.

### Tail suspension test

Mice were suspended from a horizontal bar positioned 50 cm above the floor using adhesive tape attached approximately 1 cm from the tip of the tail. To prevent tail-climbing behavior, the tail was placed through a small plastic tube. Each trial lasted for 6 min, and total immobility time was recorded using a video camera placed beside the apparatus. The video camera was connected to a computer, and behavior was tracked and analyzed in real time using ANY-maze video tracking software (Stoelting Co., Wood Dale, IL, USA). The software was set to begin scoring immobility when the mouse remained motionless for at least 1 s.

### Forced swim test

The forced swim test was performed as previously described^49^. Briefly, mice were placed individually in a plastic beaker (diameter × height = 16 × 30 cm) filled with water (24–26°C) to a depth of 14 cm. Immobility time for each animal was recorded for 10 min, and analyzed using ANY-maze software.

### RNA sequencing

RNA-seq analysis was performed using mPFC samples collected from three mice per group after completion of the behavioral tests (day 23). Total RNA was extracted from mPFC tissue using TRIzol reagent (Thermo Fisher Scientific, Waltham, MA, USA) according to the manufacturer’s instructions. The mPFC was collected from coronal brain sections using a 1.5-mm diameter tissue puncher (BrainScience Idea, Osaka, Japan) at the level of +2.0 to +1.5 mm from bregma. Libraries were prepared using the TruSeq Stranded mRNA Library Prep Kit (Illumina, San Diego, CA, USA) and sequenced on a NovaSeq 6000 platform (2 × 150 bp; Illumina) to a depth of 6 Gb per sample.

### Differential expression, interaction, and functional enrichment analyses

To reduce noise from lowly expressed genes, Transcripts Per Million (TPM)-based filtering was applied prior to differential expression analysis. For each strain (BALB/c and C57BL/6J) and experimental group (CTL and CRS; n = 3 per group), genes were classified as lowly expressed if they met either of the following criteria: (i) mean TPM < 1 across samples within a group, or (ii) TPM < 1 in at least two of the three samples within a group. Genes meeting these criteria in both the CTL and CRS groups within a given strain were designated as strain-specific exclusion targets. To retain genes expressed in at least one genetic background, only genes classified as lowly expressed in both BALB/c and C57BL/6J and mice were excluded from subsequent analyses.

For within-strain differential expression analysis, filtered count data were analyzed using the RNA-seq Analysis Portal (QIAGEN, Germantown, MD, USA). DEGs were defined as genes with an absolute log₂ fold change (|log₂FC|) > 0.263, corresponding to a fold change > 1.2 or < 0.83, and *p* < 0.05. DEGs identified in each strain were subsequently subjected to canonical pathway analysis using IPA (QIAGEN) and GO enrichment analysis using Metascape^50^.

To identify genes showing strain-dependent transcriptional responses to stress, an interaction analysis was performed using the DESeq2 package^51^ in R (v.4.6.0)^52^. A generalized linear model with the design formula ∼ strain + condition + strain:condition was fitted to the filtered count data, where the interaction term (strain:condition) was used to identify genes whose transcriptional responses to CRS differed between BALB/c and C57BL/6J mice. To preserve the original magnitude of expression changes, log_2_FC shrinkage (lfcShrink) was not applied. Interaction DEGs were defined as genes with |log₂FC| > 0.263 and *p* < 0.05. The interaction log₂FC was extracted in the direction of the CRS-induced log_2_FC in C57BL/6J mice minus that in BALB/c mice. Interaction DEGs were subsequently subjected to canonical pathway and upstream regulator analyses using IPA software.

### Statistical analysis

The normality of data distribution was assessed using the Shapiro–Wilk test. Time-course changes in body weight from day 1 to day 21 were analyzed using a two-way repeated measures ANOVA. Within-group changes in body weight between day 1 and day 21 were analyzed using a paired *t*-test. For non-normally distributed data, the non-parametric Mann–Whitney *U* test was used. All analyses were performed using GraphPad Prism v.9.0 (GraphPad Software, Inc., San Diego, CA, USA) or R software v.4.6.0. Differences were considered statistically significant when *p* < 0.05.

## Acknowledgements

We thank the staff of the Animal Facility at Saga University for their technical assistance. LC-MS/MS analysis was conducted at the Analytical Research Center for Experimental Sciences, Saga University.

## Funding

This work was supported by JSPS KAKENHI (Grant Numbers 24K17849 to T.K., 25K02370 to T.U., and JP23K16819 to I.E.), and by the Japan Science and Technology Agency Support for Pioneering Research Initiated by the Next Generation (JST SPRING; Grant Number JPMJSP2114) at Shinshu University. Research equipment used in this study was supported by the Ministry of Internal Affairs and Communications of Japan (Grant Number JPMI10001 to A.M.) at Saga University and by the MEXT Project for Promoting Public Utilization of Advanced Research Infrastructure (Program for Supporting the Construction of Core Facilities; Grant Number JPMXS0441000021) at Shinshu University.

## Author contributions

T.K., A.W.O., Y.N., and A.I. conducted the behavioral experiments. T.K., A.W.O., and T.U. prepared total RNA. T.K., A.W.O., T.S., E.K., and S.K. performed RNA experiments and IPA analyses. S.K. and M.Y. performed the statistical analyses. A.M., I.E., and G.Y. performed biochemical analyses of serum samples. T.K., A.W.O., and T.U. wrote the manuscript. M.Y. edited the manuscript. T.K. and T.U. designed the study, and H.Y. and T.U. supervised the study.

## Competing interests

The authors declare no competing interests.

## Data availability statement

The RNA-seq datasets generated in this study have been deposited in the NCBI Gene Expression Omnibus (GEO) repository under the accession number GSE330744 and are accessible at [https://www.ncbi.nlm.nih.gov/geo/query/acc.cgi?acc=GSE330744].

## Additional information

Correspondence and requests for materials should be addressed to H.Y. or T.U.

## Notes

### Competing Interest Statement

The authors have declared no competing interest.

