## Supplementary Information for "Differential physiological, behavioral, and medial prefrontal cortex transcriptomic responses to chronic restraint stress between BALB/c and C57BL/6J mice"

[1]Division of Physiology, Faculty of Medicine, Saga University, Saga 849-8501, Japan
[2]Department of Biomedical Engineering, Graduate School of Medicine, Science and Technology, Shinshu University, Nagano 390-8621, Japan
[3]Department of Orthopedic Surgery, Shinshu University School of Medicine, Nagano 390-8621, Japan
[4]Department of Social and Environmental Medicine, Saga University, Saga 849-8501, Japan
[5]The United Graduate School of Agricultural Sciences, Kagoshima University, Kagoshima 890-0065, Japan.
[6]Laboratory of Biochemistry, Faculty of Agriculture, Saga University, Saga 840-8502, Japan.
[7]Division of Gene Research, Research Center for Advanced Science and Technology, Shinshu University, Nagano, 390-8621, Japan
[8]Institute for Biomedical Sciences, Research Cluster for Social Implementation, Shinshu University, Nagano, 390-8621, Japan
[9]Department of Behavioral Physiology, Faculty of Medicine, University of Toyama, Toyama, Japan
[10]Research Center for Idling Brain Science, Life Science Research Center, University of Toyama, Toyama, Japan
[11]These authors contributed equally to this work.

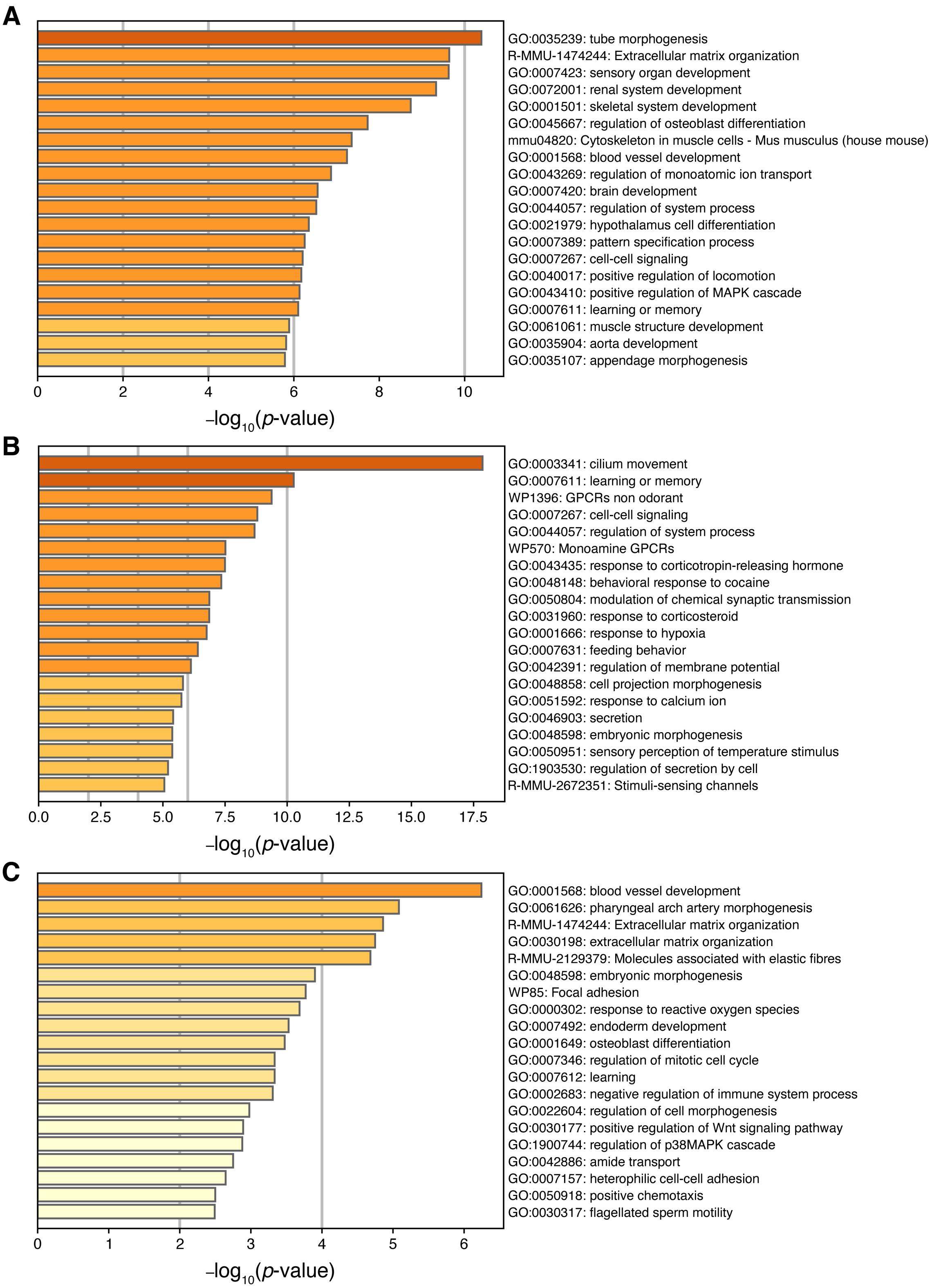


**Supplementary Figure 1**. Gene ontology and pathway enrichment analysis of differentially expressed genes in the mPFC following CRS. (A–C) Gene ontology (GO) and pathway enrichment analysis was performed using Metascape on differentially expressed genes ($|\log_2 FC| > 0.263$, $p < 0.05$) identified in each comparison. Bar graphs show the top 20 enriched terms ranked by $-\log_{10}$($p$-value). Enriched terms for differentially expressed genes (DEGs) identified in BALB/c mice (CRS vs. CTL) (A), C57BL/6J mice (CRS vs. CTL) (B), and by the strain × stress interaction analysis (C), respectively.

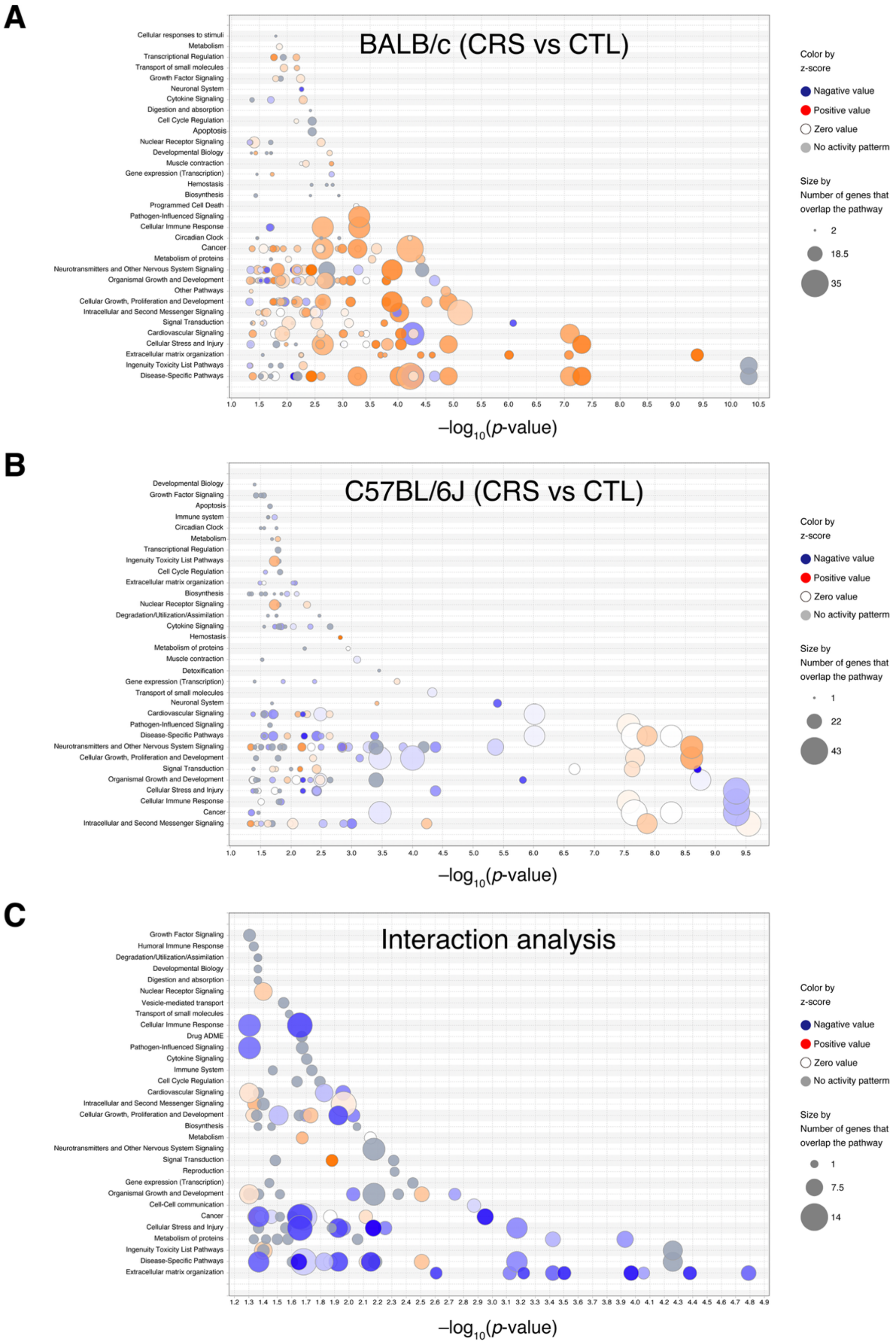


**Supplementary Figure 2**. Overview of IPA canonical pathway analysis of differentially expressed genes in the mPFC following CRS. Bubble plots summarize the results of IPA canonical pathway analysis performed on differentially expressed genes ($|log_2FC| > 0.263$, $p < 0.05$). Only pathways reaching statistical significance ($p < 0.05$) are displayed. Bubble size represents the number of DEGs overlapping with each pathway, and bubble color indicates the activation z-score (red, activated; blue, inhibited; grey, no activity pattern available). The y-axis lists IPA-defined canonical pathways. Canonical pathway analysis of DEGs identified in BALB/c mice (CRS vs. CTL) (A), in C57BL/6J mice (CRS vs. CTL) (B), by strain × stress interaction analysis (C), respectively.
